# *Cucumber Mosaic Virus* and their associated satellite RNAs infecting banana (*Musa sp. Genomic group AAA)* in Côte d’Ivoire : A molecular characterization

**DOI:** 10.1101/2020.11.17.386326

**Authors:** Dominique K. Koua, Anicet Ebou, Theodore K. Kouadio, Philippe Lepoivre, Sébastien Massart

## Abstract

In Côte d’Ivoire, banana (*Musa sp.*) ranks third among exportation products and represents 3% of the Gross Domestic Product with a national production up to 500000 tons in 2019. Banana is subject to numerous disease agents among which viruses cause significant losses. To figure out the impact of viruses in Ivorian industrial banana fields, surveys were conducted in the 7 main banana production departments. A total of 260 leaf fragments presenting viral symptoms were collected and analyzed. From the 65 leaf fragments used for biological indexing, 14 showed symptoms related to *Cucumber mosaic virus* (CMV). CMV presence was confirmed by double-antibody sandwich enzyme-linked immunosorbent assay (DAS-ELISA) using CMV polyclonal antibodies. CMV strains we isolated, appeared to be highly infectious and to produce various symptoms like mosaic, chlorosis, and necrotic spots on *Cucumis sativus*, *Cucurbita pepo*, and *Nicotiana tabacum*. Satellite RNAs (SatRNAs) associated with CMV isolates were also detected using reverse transcription polymerase chain reaction (RT-PCR) with a degenerate primer pair. CMV’s coat protein as well as satRNAs was sequenced. Novel Ivorian coat proteins and satRNAs were compared to publicly available sequences. We noticed a single amino acid substitution (Serine to Leucine) at position 73 of the novel coat protein that allowed us to divide Ivorian CMV strains into two groups. Molecular and phylogenetic analysis suggested that Ivorian strains might be classified into CMV Subgroup IA. We also discovered that satellite RNA associated with Ivorian CMVs form a separate clade.

## Introduction

Côte d’Ivoire is the first African exporter of banana to European countries [1]. Banana represents 3% of Gross Domestic Product (GDP) of the country. The banana sector represents 9,000 direct jobs and 35,000 indirect jobs in Côte d’Ivoire. With a production of nearly 450,000 tonnes of bananas in 2019, Ivory Coast is at the forefront of African producing countries, and records a turnover of 145 billion CFA francs [2]. Banana is therefore of importance for the national economy of Côte d’Ivoire.

*Cucumber mosaic virus* (CMV) is one of the top ten plant virus in molecular plant pathology and is reported to infect more than 1200 plant species in over 100 families, both monocots and eudicots: fruit crops, vegetables, and ornamental, including banana [3]. CMV belongs to the genus *Cucumovirus* and the family of *Bromoviridae.* CMV is a positive sense RNA plant virus with a tripartite genome [4–6]. This genome encodes five proteins: the 1a and 2a proteins are encoded by RNA1 and RNA2 respectively and are obligatory for viral replication. The 2b protein is encoded by a subgenomic RNA (RNA4A) from RNA2, and functions as a viral suppressor of RNA silencing [7]. RNA3 is bicistronic: the protein encoded by the 5′ open reading frame (ORF) is the designated movement protein, while the coat protein (CP) encoded by the 3′ ORF is translated from a subgenomic RNA4 synthesized de novo from RNA3 minus-strand progeny [8]. CMV strains are broadly divided into two subgroups (I and II) based on nucleotide sequence homology. CMV strains in the same subgroup share a high degree of sequence similarity [9]. Several CMV strains discovered on banana from Latin America have already been described by proteomics and phylogenetics [5,8–12] but very few analyses has been carried out on strains originating from West Africa. Although CMV has been well studied biologically [13] and from a molecular point of view in Asian and American countries, molecular and phylogenetic characterization of African CMV strains targeting banana crops is still missing.

CMV is not known as a major virus infecting banana but causes notable losses and has been reported as one of the virus infecting banana in Cote d’Ivoire [12,13].

In this study, we report one of the first molecular and phylogenic based characterization of Ivorian CMV strains identified on banana Cavendish, cvs Great Naine, and William, AAA group. These results are based on the analysis of an exhaustive survey that led to the collection of 260 banana leaf fragments from symptomatic banana plants in the 7 main banana production areas of Côte d’Ivoire. After antibody-based virus detection, 14 leaf samples were used for PCR amplification and DNA sequencing while 65 samples were used for biological indexing. In addition to classical biological comparison (description of induced symptoms on reporting plants) and nucleotide comparison of coat protein, we sequenced partially the detected micro-satellite of 15 CMV isolates.

## Materials and methods

### Sample collection and storage

Banana (variety Cavendish, cvs Great Naine and William, AAA group) leaf samples showing symptoms of possible viral etiology were collected in the seven major production areas of culture from 2010 to 2011. Sampling places are located in the South and South-East of the country and correspond to banana production area around the main harbor, Abidjan (Fig 1). A total of 260 bananas leaves were collected in industrial plantations. A sample consisted of fragments of a leaf demonstrating one or more symptomatic traits. On the field, samples were placed in a plastic bag, assigned a code, and stored in dried ice. Laboratory storage conditions consisted of a refrigerator at −20°C. A second part of each sample was kept desiccated on calcium chloride.

**Fig 1.**
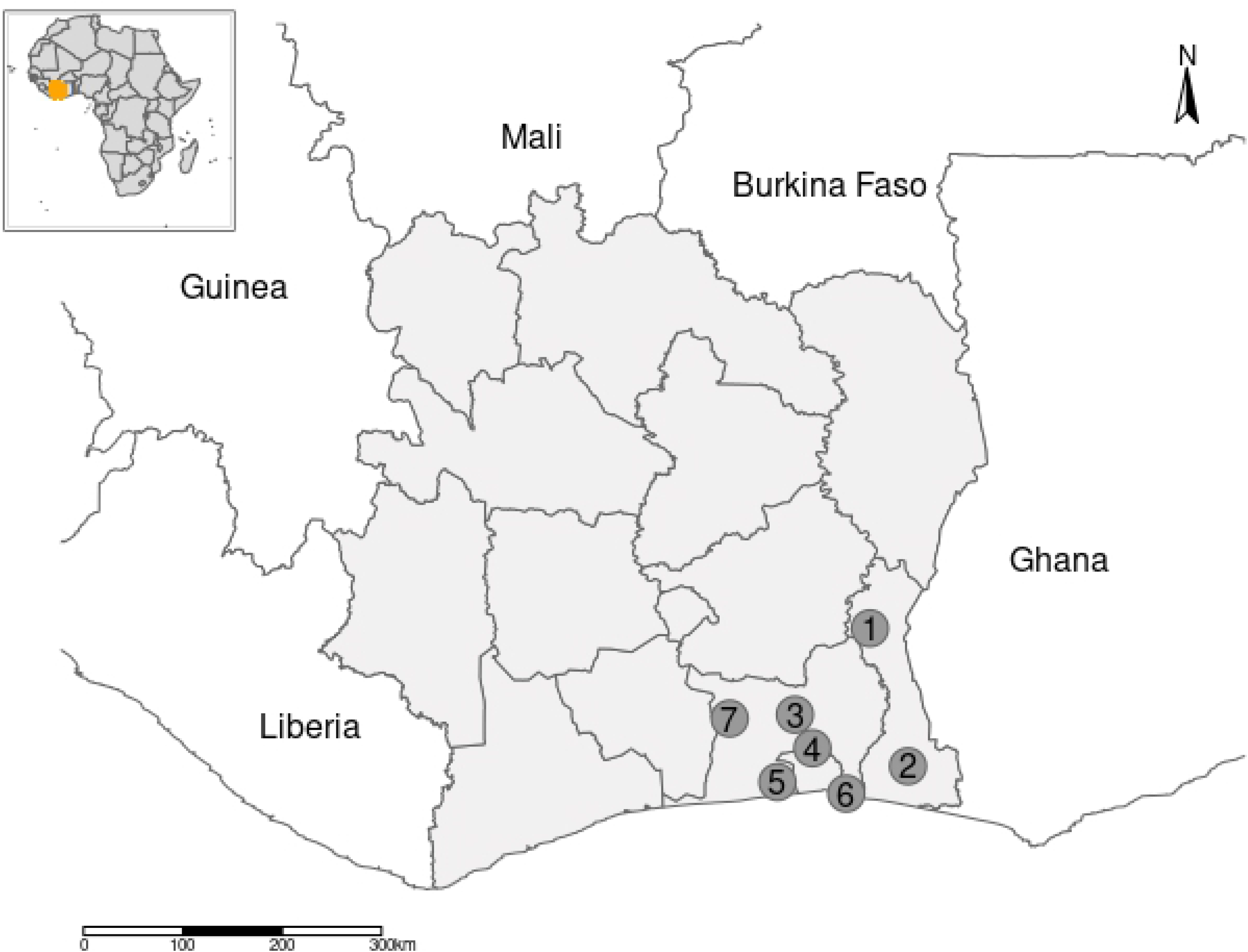
Location of sample collection areas in industrial banana plantations in Côte d’Ivoire. 1. Abengourou (Lat: 6.7146399, Long: −3.5654096), 2. Aboisso (Lat: 5.4675302, Long: −3.2185867), 3. Agboville (Lat: 5.9342509, Long: −4.2500361), 4. Azaguié (Lat: 5.6305067, Long: −4.0907336), 5. Dabou (Lat: 5.3242556, Long: −4.4098582), 6. Grand-Bassam (Lat: 5.2198516, Long: −3.7821731), 7. Tiassalé (Lat: 5.8985662, Long: −4.8458078)

### Virus detection and sequencing

CMV detection was performed using three complementary approaches: double antibody sandwich enzyme-linked immunosorbent assay (DAS-ELISA), biological indexing, and reverse transcription polymerase chain reaction (RT-PCR).

DAS-ELISA was carried out using the Loewe Phytodiagnostica GmbH kit (Sauerlach, Germany). The kit contains two polyclonal antibodies with their related conjugates that are coupled to alkaline phosphatase and also includes positive and negative probes as well as sensitization, extraction and cleansing buffers. These antibodies (anti-CMV) can detect the two groups of CMV. Leaf samples, stored at −20°C, were defrosted and crushed with a manual ball extractor (AGDIA, France). The grinding was done with a ratio of 0.5 g of sample per 5 ml of the extraction buffer. Settled and clarified juice (kept on ice) was then collected in tubes for antibody detection. Antibody detection was conducted following the manufacturer’s recommendation. Optic density was determined using a ELISA plate reader (Titertek Multiskan) after 1 and 2 hours of incubation in darkness. A sample was considered positive when the absorbance (at 405 nm) was greater than 2 times the average of absorbance of reference samples (false positive and true negative) [14,15].

Molecular detection of CMV using RT-PCR was carried out using specific primers CMV 3′ : TTTTAGCCGTAAGCTGGATGGACAACCC and CMV 5′ : TATGATAAGAAGCTTGTTTCGCGCA [16,17].

One milliliter of extraction buffer (137 mM NaCl; 8 mM Na 2 HPO 4; 1.5 mM KH 2 PO 4; 2.7 mM KCl; 80 mM Na 2 SO 3; 3 mM NaN 3; 0.05% Tween 20 – pH 7.2) was added in a grinding bag to 50 mg of dried leaves or 2 ml added to 0.5 g of fresh leaves. After 15 minutes of soaking the dried samples or immediately for fresh leaves, samples were crushed with an electric ball mill (Power Plus X022). Settle clarified juice was diluted 100 times with sterile distilled water. For RT-PCR amplification of nucleic acid sequences, reactions were made in volumes of 25 μl (20 μl of the reaction mixture + 5 μl of the sample corresponding to diluted crude extract) using the Titan RT-PCR kit (Rocher, Mannheim, Germany). A reaction mixture was composed of 5 μl of Buffer (5x Concentrate) Titan RT-PCR, 0.5 μl of dNTP, 0.5 μl CMV 3′ (25 μM), 0.5 μl CMV 5′ (25 μM), 1.25 μl of DTT (100 mM), 0.5 μl of Titan mixed enzyme, and 11.75 μl of sterile distilled water, i. e. 20 μl by mix. All amplification reactions included both negative and positive controls from banana plants grown in greenhouses as well as blanco made up of sterile distilled water. Amplification cycles was performed using a My Cycler™ (Biorad, USA) following this program: 50°C for 30min, 94°C for 5min, 40 cycles of 94°C for 30sec, 54°C for 60sec, 72°C for 2min then 72°C for 10min. The last step consisted in the revelation of amplification products on agarose gel 1% (1 g agarose per 100 ml of buffer Tris, Acetate, EDTA (TAE) 1 x) containing 10 μl of Ethidium bromide (BET), in TAE buffer 1 x. Electrophoresis migration was carried out under a constant current of 120 mA for 45 min. The gel was then visualized by UV lighting allowing the observation of amplified bands that were pictured using a digital camera.

CMV’s satellite RNAs (satRNAs) identification was carried out by amplification of nucleotide sequences by RT-PCR following the procedure described above but using primers proposed by Gafny et al. [18] and Nouri et al. [19]. These primers include header and tail sequences found in all known CMV satRNA and deposited at the National Center for Biotechnology and Information (NCBI) Genbank. A positive control, CMV strain with satellite RNA (DSMZ collection: PV-0029 and PV-0092, Germany) and a negative control (CMV strain without satellite RNA) were used. PCR products were purified with the Qiaquick PCR Purification kit (Qiagen, Benelux) and sent for sequencing using the Sanger dideoxy sequencing technology (MACROGEN Inc., Netherland). In all, 16 CMV isolates from the 7 sampling locations were sequenced from amplicons of genes partially coding for the coat protein and part of an untranslated region (UTR) of the RNA3. For the sequencing of satellite RNA, 15 RNA satellite isolates were selected from amplification and purification results.

### Indicator plant inoculation

The biological indexing was carried out by mechanic infection of *Cucumis sativus* cv Poinsett Pepino, *Cucurbita pepo* Linn. cv Precose Maraîchère, and *Nicotiana tabacum* cv. Samsun. Three test plants were used per virus isolate. Mechanical inoculation took place at the two-leaf cotyledon stage for Cucurbits (cucumber, courgette) and at the 4-6 leaf stage for tobacco plants. Banana leaf samples were ground in a bag containing phosphate buffer at pH 7.2 (KH 2 PO 4 0.05 M; DIECA 0.01 M) in a ratio of 1:5 (g:ml) for samples stored at −20°C and 1:10 (g: ml) for dried samples. The shred was clarified and collected in a Petri dish, then a pinch of carborundum and a pinch of activated carbon were added. After wearing gloves to avoid any contamination, healthy plant leaves previously sprinkled with carborundum were inoculated by rubbing their top face with a finger moistened in the inoculum. The inoculated leaves were immediately rinsed with distilled water and inoculated plants were shade-stored in greenhouse for 24 hours: (25°C, 16 hours of auxiliary lighting at the phytopathology laboratory, Gembloux) or in anti-insect cages (ambient temperature average of 30°C at the phytopathology laboratory, INP-HB). Two CMV isolates from the DSMZ collection (DeutscheSammlung Von Mikroorganismen und Zell-kulturen, PV-0029 and PV-0092; Germany) and healthy plants were used as positive and negative controls respectively. All plants (inoculated or not) were regularly watered and monitored during 30-35 days.

### Sequence analyses

Data issued from sequencing were visualized using multiple sequence alignment obtained from Clustal Omega [20]. Similarity searches were realized using the BLASTN program from BLAST suite [21] and using NCBI GenBank non redundant nucleotide database as the search database (downloaded and compiled in December 2019). Phylogenetic trees were built using the maximum-likelihood method with 1000 bootstrap replicates under the MEGA 7 software [22]. A data set containing 49 sequences from the different CMV subgroups (S1 Table) and 1 sequence used as the outgroup (ER-PSV: U15730) were used to build CMVs coat proteins tree. A data set of 50 sequences from known CMV’s satRNAs (S2 Table) was used for the satRNAs tree. Phylogenetic trees were visualized using iTOL v4 [23].

## Results

### Symptoms observation and CMV detection

Virus presence was detected by measuring the absorbance of a solution of clarified leaf juice incubated with antibodies (DAS-ELISA). Absorbance measured for negative control used during our experiment varied between 0.09 and 0.12. For the positives controls, absorbance ranged from 2.65 to 3.32. Absorbance measured for studied samples ranged from 0.10 to 3.47. A total of 248 samples out of 260 were tested positive for CMV with absorbance ranging from 0.18 to 0.24. DAS-ELISA realized on symptomatic leafs therefore allowed to confirm the presence of CMV: 85.7% of the sample from Abengourou (East) were positive to CMV and 100% for samples from Aboisso (south-East). Table 1 summarizes, for each sample place, percentages of symptomatic banana samples that appeared to be positive to CMV.

**Table 1.**
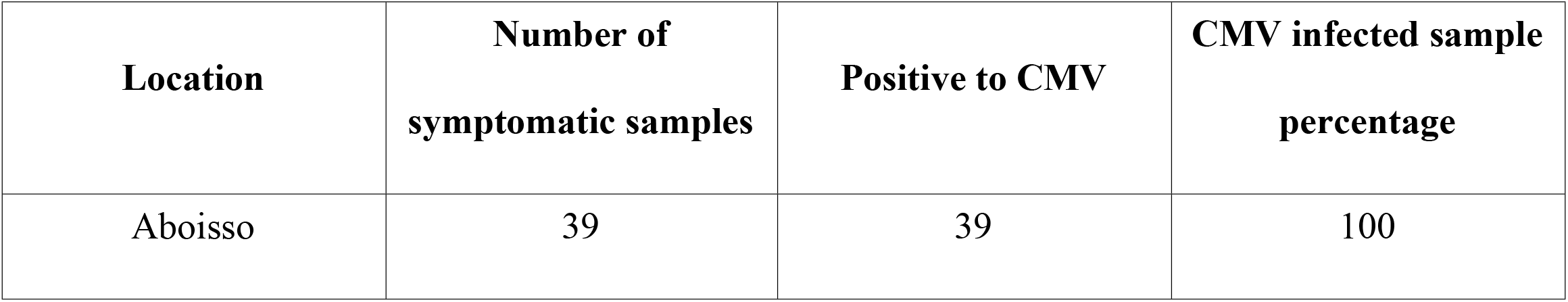
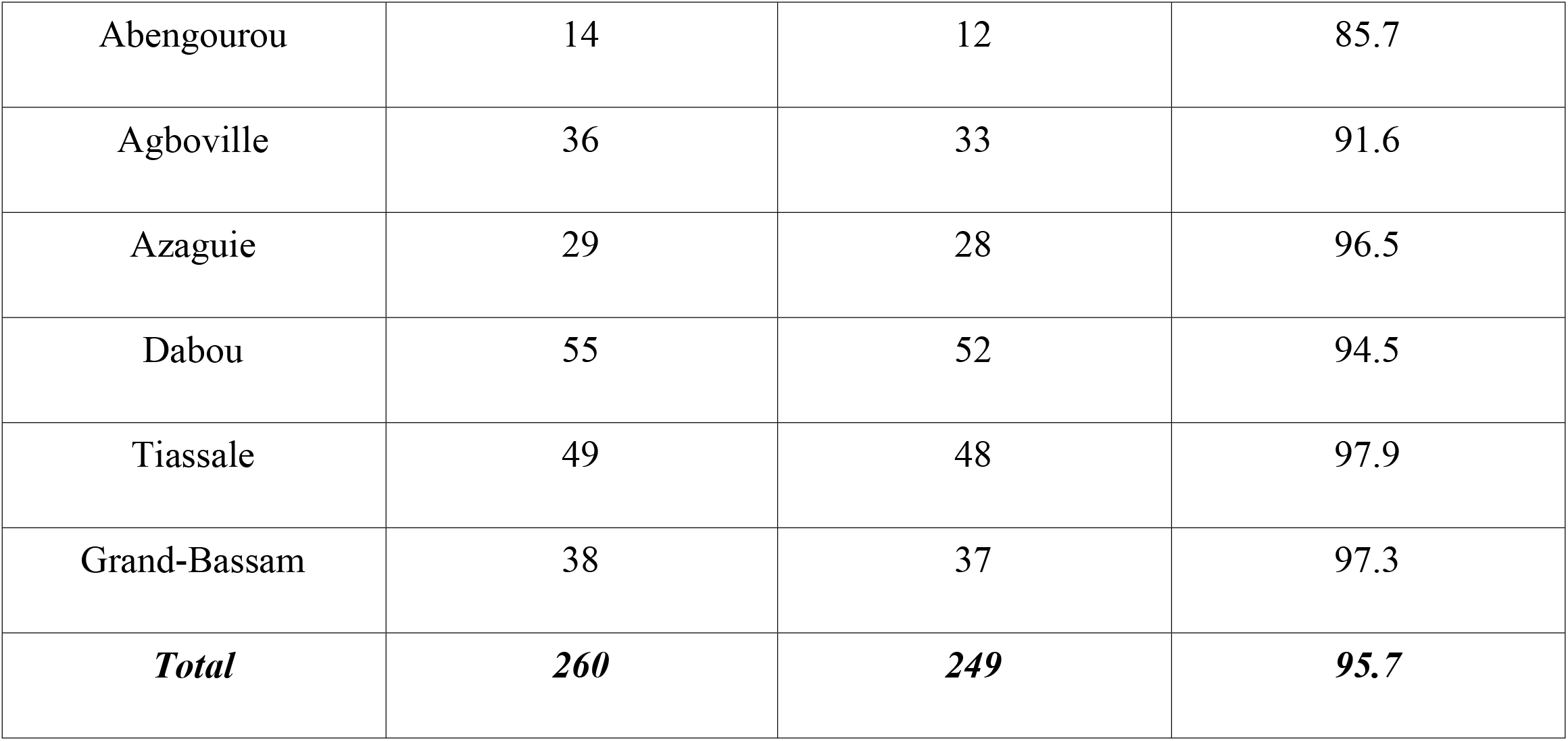
Percentages of symptomatic banana samples positive to CMV, by sampling sites.

The most common symptoms observed countrywide are chlorosis, mosaics and leaf necrosis. Observations concerning the main symptoms associated with banana leafs samples positive to CMV are provided in Table 2.

**Table 2.**
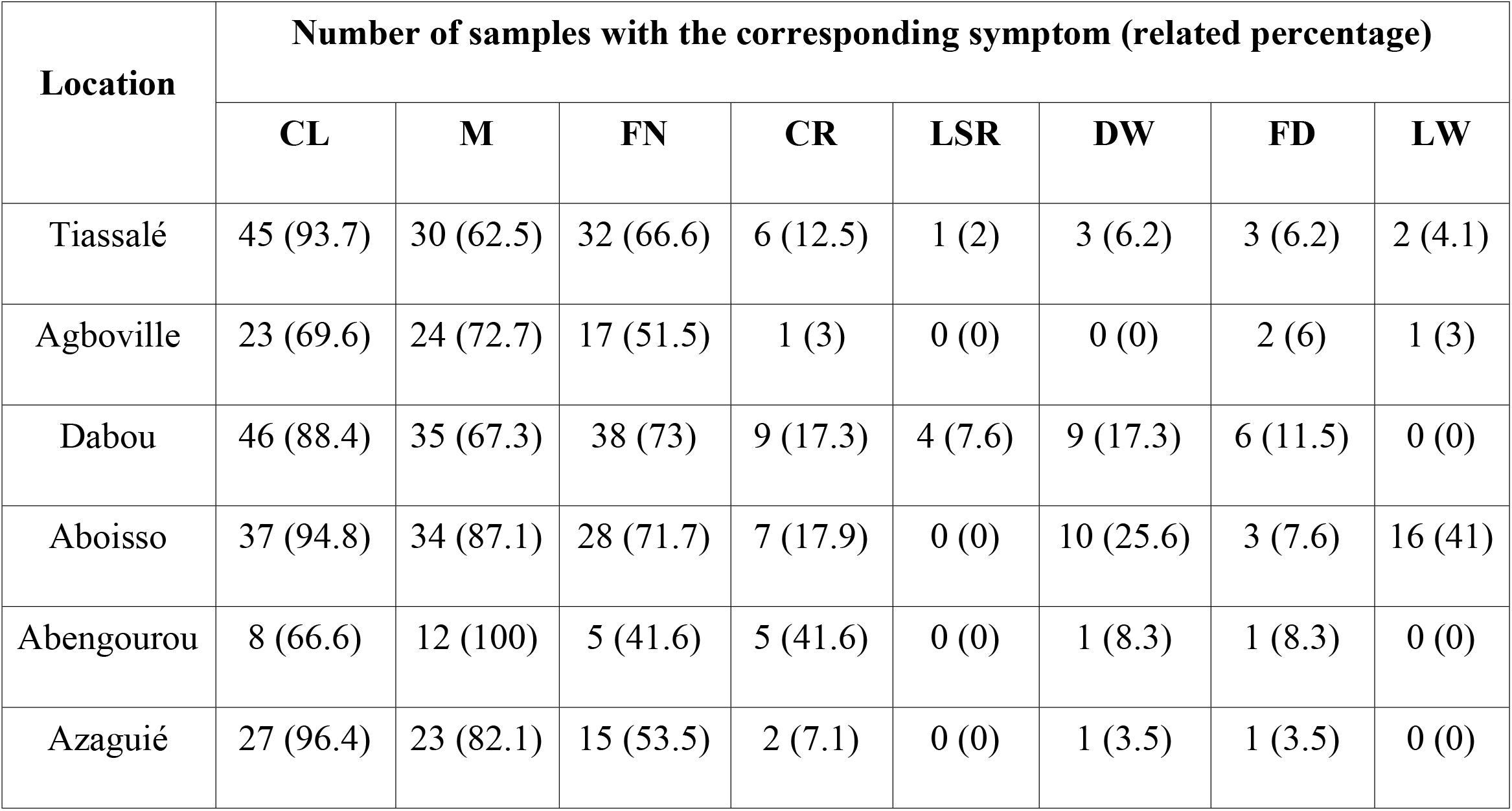
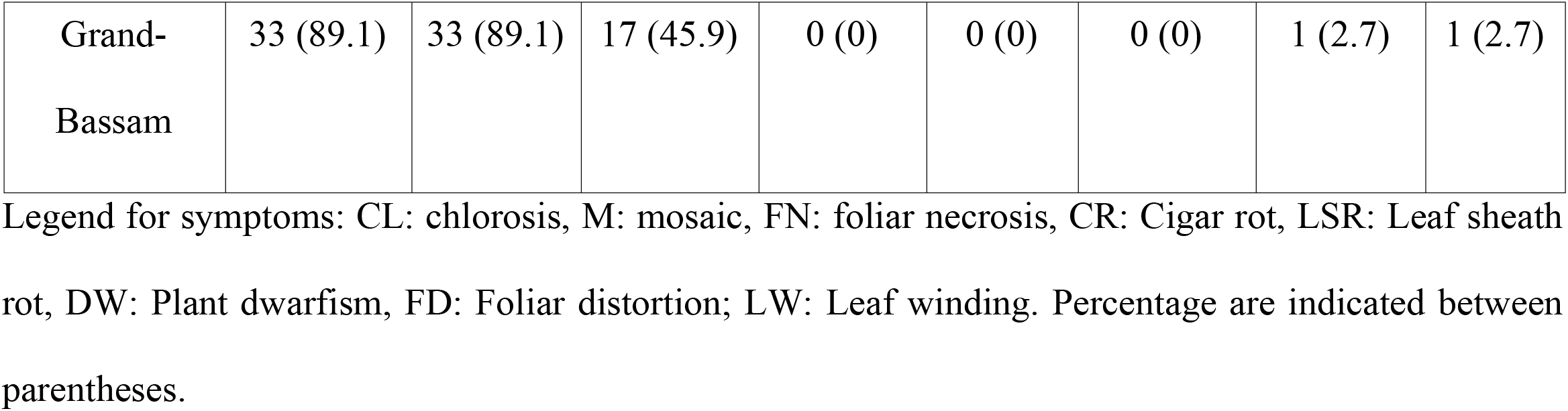
Symptoms associated with samples positive to CMV for each collection place.

Fragments of expected size (950 bp) were identified in infected samples along with positives control (Fig 2). RT-PCR test confirmed the serologic test for all but the BT16 sample that finally appeared positive even if the ELISA test was negative. These results indicate that CMV was effectively present in 249 samples collected in the 7 areas known to be major production places of banana in the country.

**Fig 2.**
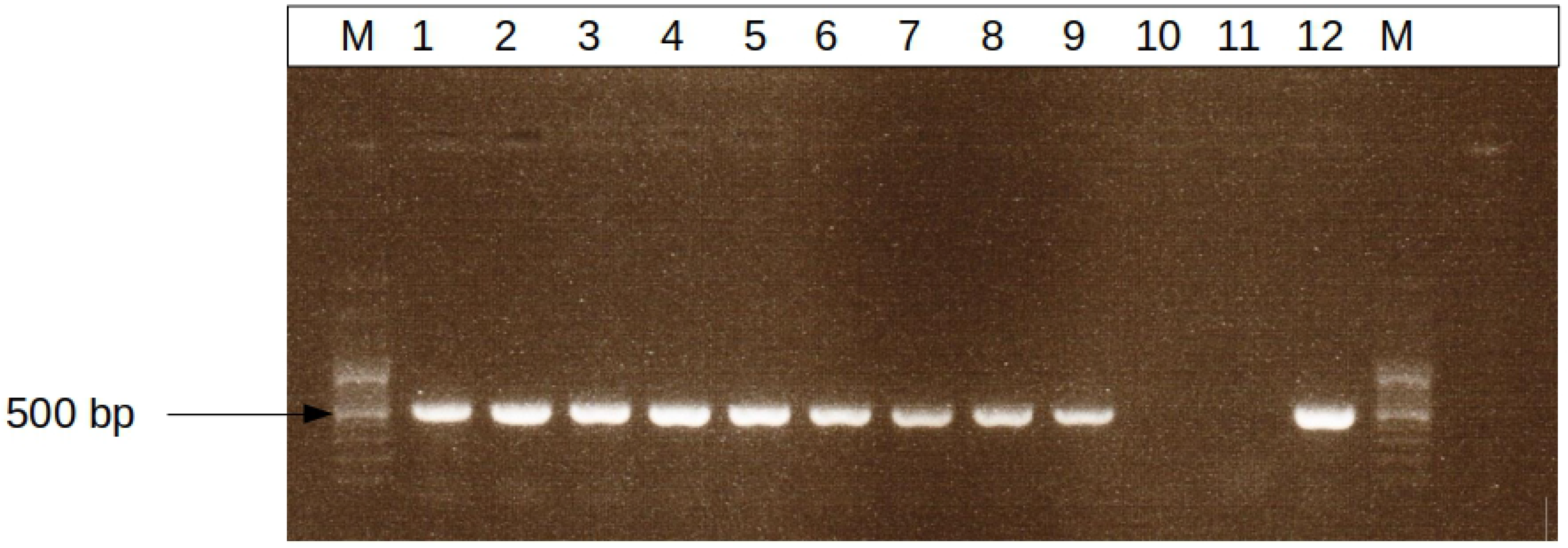
Agarose gel electrophoresis of CMV’s RT-PCR. M=100 pb (Ladder DNA) molecular weight marker, 1 to 9: 500pb amplicons from 9 symptomatic banana samples; 10: mix + sterile distilled water, 11: healthy plant in greenhouse, 12: CMV positive control.

### Symptomatology on indicator plants

The symptom development on indicator plants was evaluated for CMV isolates. The positive controls caused symptoms of local chlorotic lesions 3-4 days after inoculation and yellow mosaics 8 days after inoculation on tobacco plants.

A total of 14 samples out of the 65 tested induced symptoms on the indicator plants. These symptoms included chlorosis and/or mosaic associated with leaf puffiness (S3 Table). Symptoms appeared 4 to 5 days after inoculation on Cucurbits and 14 days after inoculation for tobacco plants. The following isolates produced symptoms: AZ1, AZ2, AZ3, AZ5, AZ13 from Azaguié; BM32 and BM40 from Grand-Bassam; AG6, AG10, AG20, AG24 from Agboville; BT18 from Tiassalé; SP24 from Dabou and CA26 from Aboisso. Fig 3 illustrates some of the observed symptoms on reporting plants (zucchini, cucumber, and tobacco).

**Fig 3.**
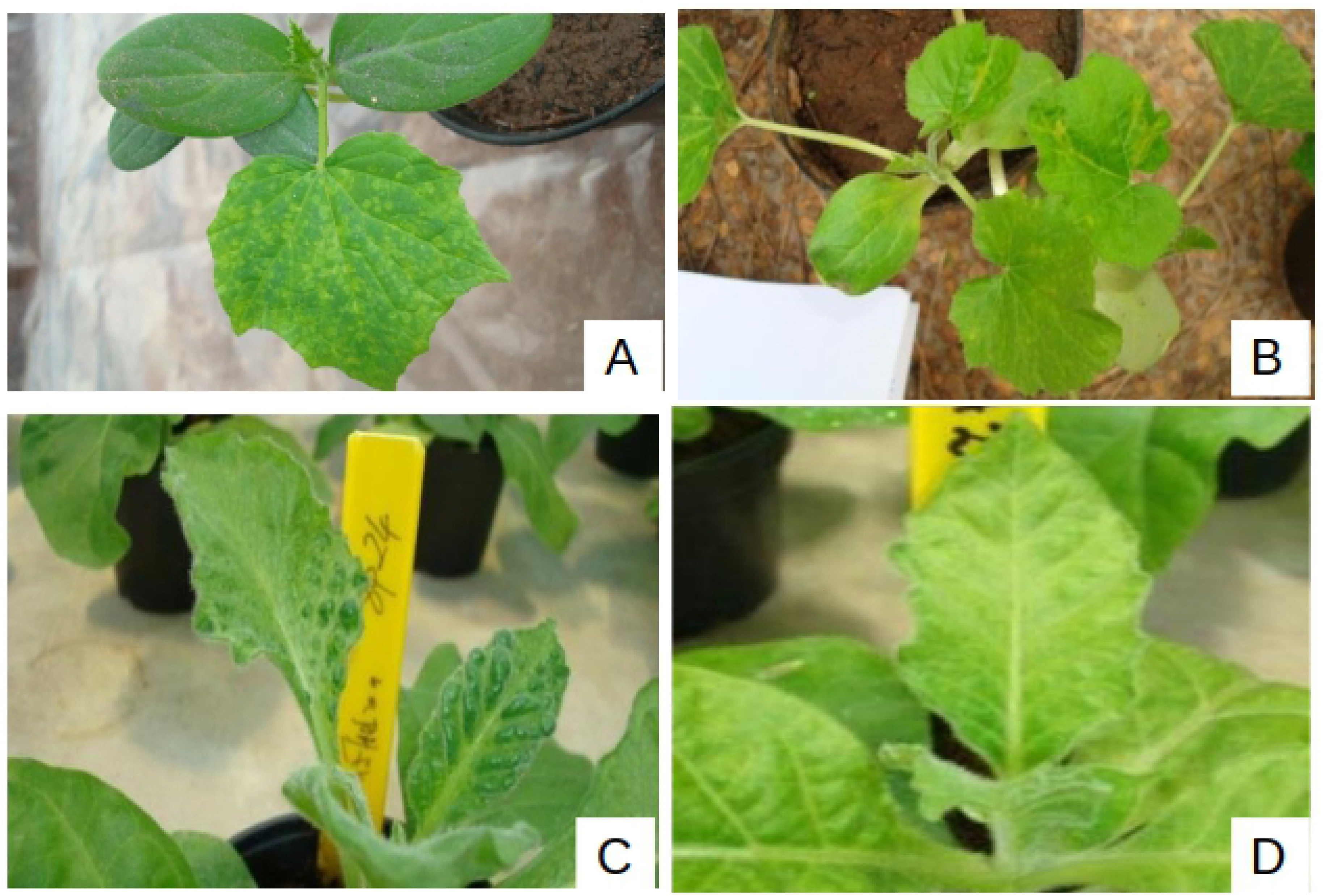
Symptoms observed on reporting plants after CMV’s biological indexing. Top-right: Mosaic on a cucumber leaf inoculated with AZ1 isolate, 7 days after inoculation; Top-left: Chlorosis on Zucchini leaves inoculated with AZ2 isolate, observed 12 days after inoculation; Bottom-right: Chlorosis including blisters on leaves of *Nicotiana tabacum* inoculated with isolate SP24, observed 20 days after inoculation; Bottom-left: Mosaic on leaves of *Nicotiana tabacum* inoculated by isolate CA26, 20 days after inoculation.

### Analysis of nucleotide sequence of CMV coat proteins

Amplicons including a portion of the coat protein and the RNA3 3’-UTR of 16 samples were sequenced and resulted in 11 unique sequences that were Further analyzed. Similarity scores are reported in Table 3.

**Table 3:** Similarity matrix between sequences of RNA3 3’-UTR of Ivorian CMV strainsa.

|  | **BM26** | **AZ6** | **AB15** | **ST17** | **AG6** | **ST18** | **AB12** | **BM32** | **AZ2** | **AG10** | **BT18** |
| --- | --- | --- | --- | --- | --- | --- | --- | --- | --- | --- | --- |
| **BM26** | 100 | 98,9 | 97,3 | 98,1 | 97,5 | 98,1 | 97,3 | 96,1 | 97,9 | 98,1 | 96,9 |
| **AZ6** |  | 100 | 98,3 | 98,1 | 98,1 | 99,1 | 96,9 | 96,9 | 98,3 | 98,7 | 97,7 |
| **AB15** |  |  | 100 | 96,9 | 97,1 | 97,9 | 96,7 | 97,3 | 97,3 | 97,7 | 97,1 |
| **ST17** |  |  |  | 100 | 97,7 | 98,3 | 96,7 | 96,7 | 96,7 | 97,1 | 96,3 |
| **AG6** |  |  |  |  | 100 | 98,1 | 96,5 | 96,5 | 97,5 | 97,1 | 98,9 |
| **ST18** |  |  |  |  |  | 100 | 97,1 | 96,5 | 97,7 | 97,7 | 98,1 |
| **AB12** |  |  |  |  |  |  | 100 | 97,3 | 96,1 | 96,7 | 96,3 |
| **BM32** |  |  |  |  |  |  |  | 100 | 96,1 | 96,3 | 95,9 |
| **AZ2** |  |  |  |  |  |  |  |  | 100 | 98,1 | 97,5 |
| **AG10** |  |  |  |  |  |  |  |  |  | 100 | 97,1 |
| **BT18** |  |  |  |  |  |  |  |  |  |  | 100 |

It appeared that Ivorian strains share a similarity of at least 94,3%. The Basic Local Alignment (BLAST) analysis indicated that sequences we identified in this study share a similarity of 94% to 98% with CMV belonging to the CMV Subgroup IA. The tree resulting from the multiple sequence alignment is provided in Fig 4.

**Fig 4.**
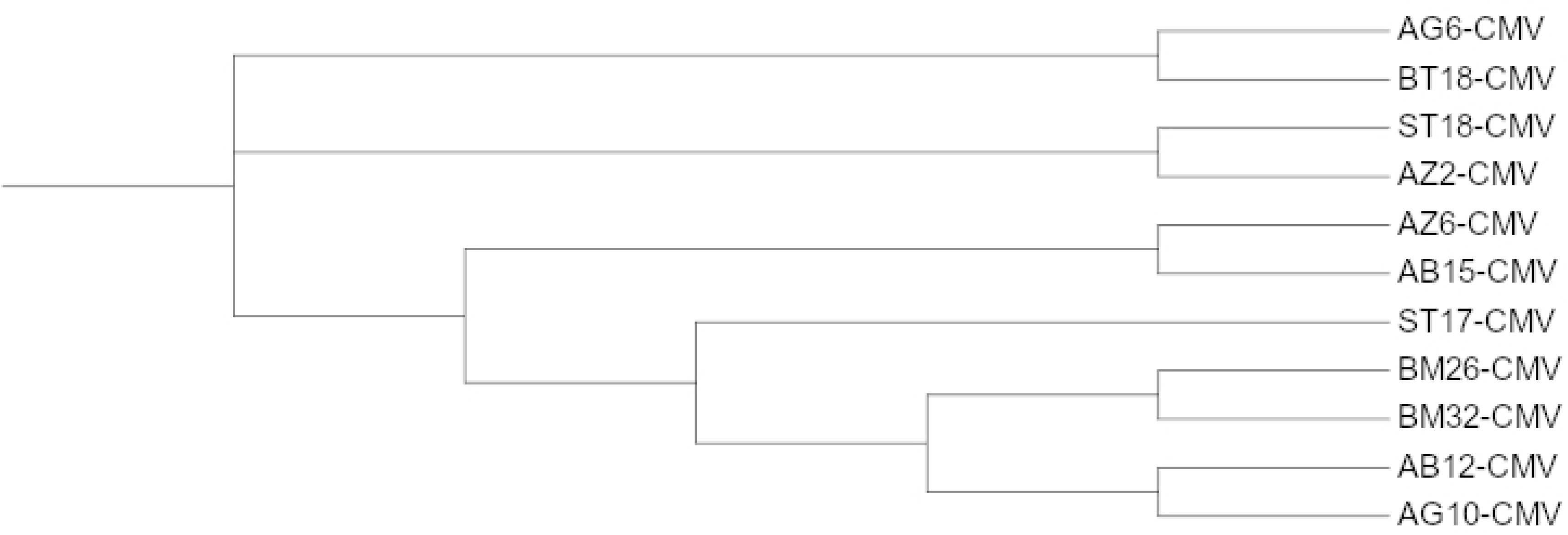
CMV’s strains phylogenetic tree using Maximum Likelihood method and JTT matrix-based model. The tree with the highest log likelihood is shown. A uniform rate is used among sites. The bootstrap consensus tree inferred from 1000 replicates is taken to represent the evolutionary history of the taxa we analyzed. Branches corresponding to partitions reproduced in less than 50% bootstrap replicates are collapsed.

The 11 distinct coat protein sequences were deposited on GenBank with accession numbers ranging from KC189911 to KC189921 (Table 4).

**Table 4.** List of identifiers of coat protein and satRNA sequence submitted to the NCBI GenBank Database.

| **Strains reference** | **Collection place** | **GenBank identifier of**<br>**Coat Protein** | **GenBank identifier of**<br>**Satellite RNA** |
| --- | --- | --- | --- |
| BM26 | Grand-Bassam | KC189911 | KC713593 |
| AZ6 | Azaguié | KC189912 | MT968978 |
| AZ15 | Azaguié | KC189912 | MT968984 |
| AB15 | Abengourou | KC189913 | MT968985 |
| ST17 | Tiassalé | KC189914 | MT968977 |
| AG6 / AG16 | Agboville | KC189915 | MT968975 |
| ST18 | Tiassalé | KC189916 |  |
| AB12 | Abengourou | KC189917 |  |
| BM13/ BM32 | Grand-Bassam | KC189918 | MT968976 |
| BM15 | Grand-Bassam | KC189918 | MT968983 |
| AZ2 / AZ36 | Azaguié | KC189919 | MT968979 |
| AG10/ AG33 | Agboville | KC189920 | MT968980 |
| BT18 | Tiassalé | KC189921 |  |
| SP16 | Dabou | KC189919 | MT968986 |
| SP24 | Dabou | KC189919 | MT968988 |
| SP30 | Dabou | KC189919 | MT968982 |
| SP37 | Dabou | KC189919 | MT968981 |
| SP39 | Dabou | KC189919 | MT968987 |

Multiple sequence alignment of Ivorian CMV coat protein sequences revealed a sequence similarity of at least 99%. The only difference between our sequences concerns position 73. We noticed an amino acid substitution: Serine to Leucine at position 73. The alignment of identified coat protein sequences is provided as Fig 5.

**Fig 5.**
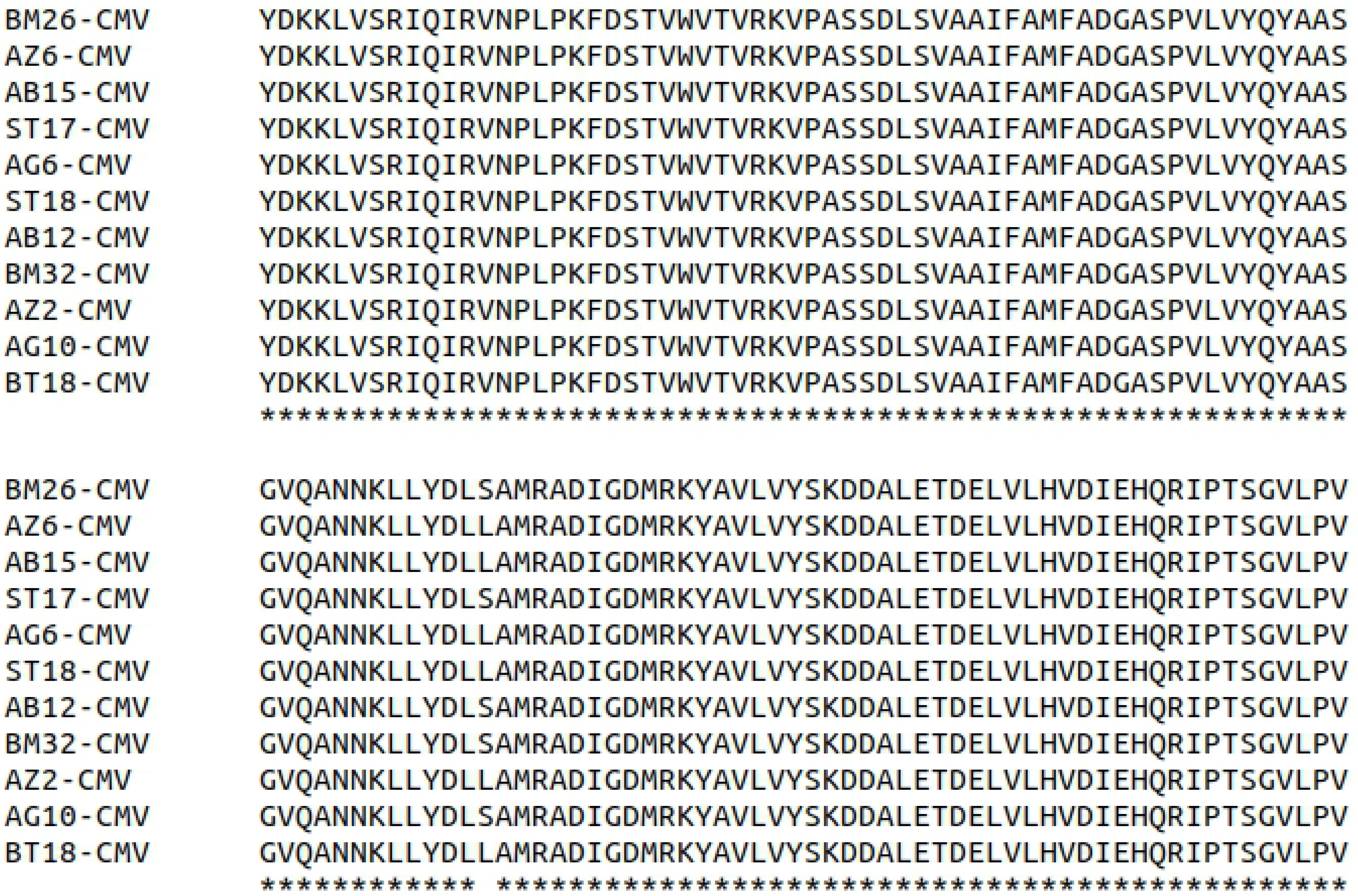
Multiple sequences alignment of Ivorian CMV strains.

Phylogenetic analysis combining Ivorian and worldwide CMV’s strains showed KC189912 (AZ6) and KC189911 (BM26) sequences to be closely related to FC-CMV and BAN-CMV respectively. FC-CMV is originated from Israel and BAN-CMV was identified in United States (Fig 6).

**Fig 6.**
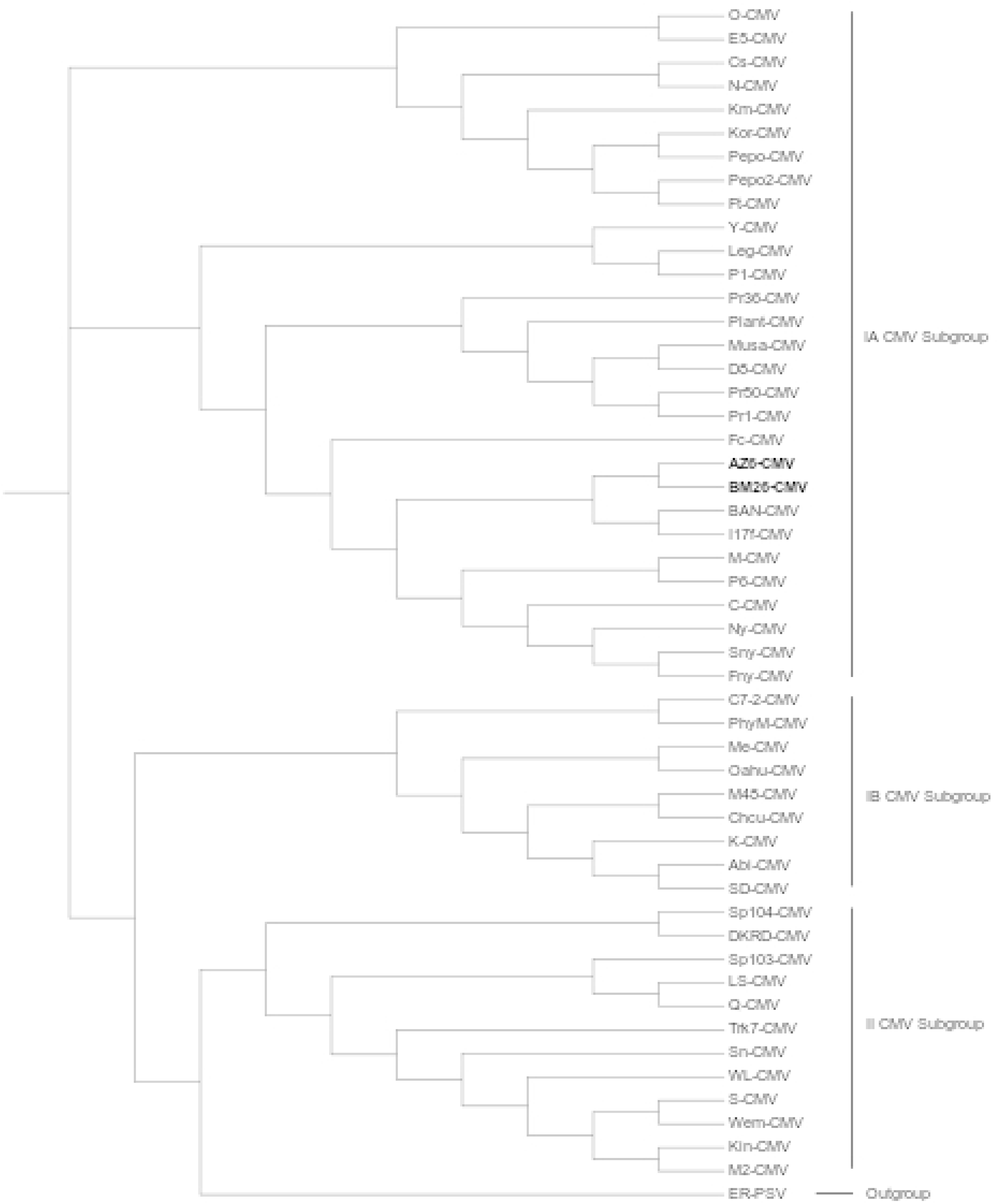
Estimation of Ivorian CMV’s phylogenetic relationship with known CMVs using the Maximum Likelihood method and JTT matrix-based model. Ivorian strains are in bold. The tree with the highest log likelihood is shown. A uniform rate is used among sites. The bootstrap consensus tree inferred from 1000 replicates is taken to represent the evolutionary history of the taxa analyzed. Branches corresponding to partitions reproduced in less than 50% bootstrap replicates are collapsed.

### Analysis of satellite RNA associated with Ivorian CMVs

RT-PCR tests revealed the presence of satellite RNA (satRNAs) in 35 strains among the 249 isolates confirmed to present CMV (Fig 7). SatRNA occurrence in this stuudy is about 14%

**Fig 7.**
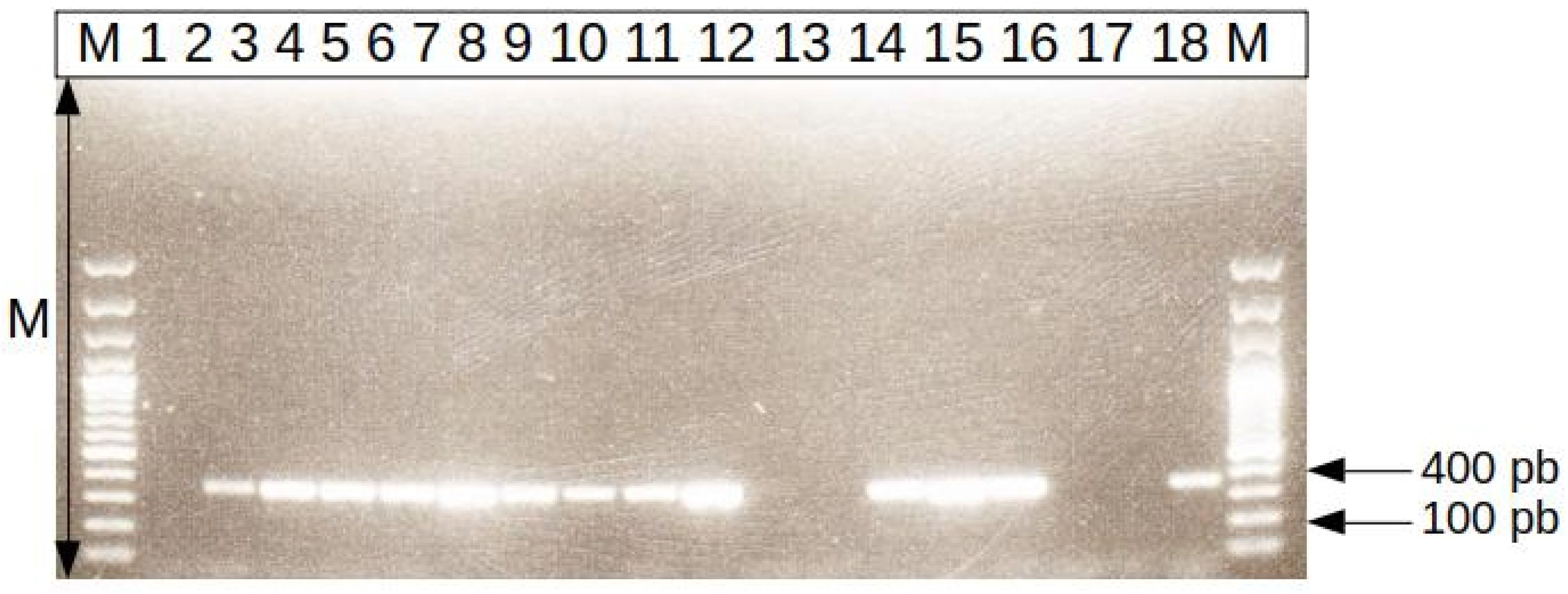
Agarose gel electrophoresis of amplification products obtained by RT-PCR for the detection CMV satellite RNAs. M : molecular mass marker (weight 100 pb) (Ladder DNA), 1 : CMV isolate without satRNA, 2 to 10 and 13 to 15: amplicons (about 350 pb) of 12 banana samples; 11 : mix+sterile distilled water, 12 : healthy plant from greenhouse, 16 to 17: CMV negative controls without satARN; 18: CMV positive control with satellite RNA (DSMZ).

At least one strain with satRNAs was detected in each sampling place except the locality of Aboisso (Table 5).

**Table 5.** Proportions of CMV satellite RNAs detected in banana plantations.

| Location | Number of symptomatic samples | Positive to CMV | Rate of infected | Number of isolates with RNA satellite |
| --- | --- | --- | --- | --- |
| Aboisso | 39 | 39 | 100 | 0 |
| Abengourou | 14 | 12 | 85.7 | 2 |
| Agboville | 36 | 33 | 91.6 | 3 |
| Azaguie | 29 | 28 | 96.5 | 10 |
| Dabou | 55 | 52 | 94.5 | 12 |
| Tiassale | 49 | 48 | 97.9 | 6 |
| Grand-Bassam | 38 | 37 | 97.3 | 2 |
| **Total** | **260** | **249** | **95.7** | **35** |

Five samples (AZ1, AZ13, BM40, BT18, SP24) out of the fourteen that induced symptoms on reporting plants appeared to possess a satRNA.

The sequencing of satRNA amplicons resulted in 15 unique sequences of 314 to 321 nucleotide long. Ivorian satRNA sequences were deposited on Genbank and are available under accession numbers ranging from MT968975 to MT968988 and KC713593 (Table 4). Fragment of satRNA identified in this study share 93.04% to 96.52% similarity.

We compute a tree with satRNA associated with Ivorian CMV and previously deposited satRNA sequences downloaded from Genbank. It appears that Ivorian satRNAs sequences clearly form a distinct clade (Fig 9).

**Fig 8.**
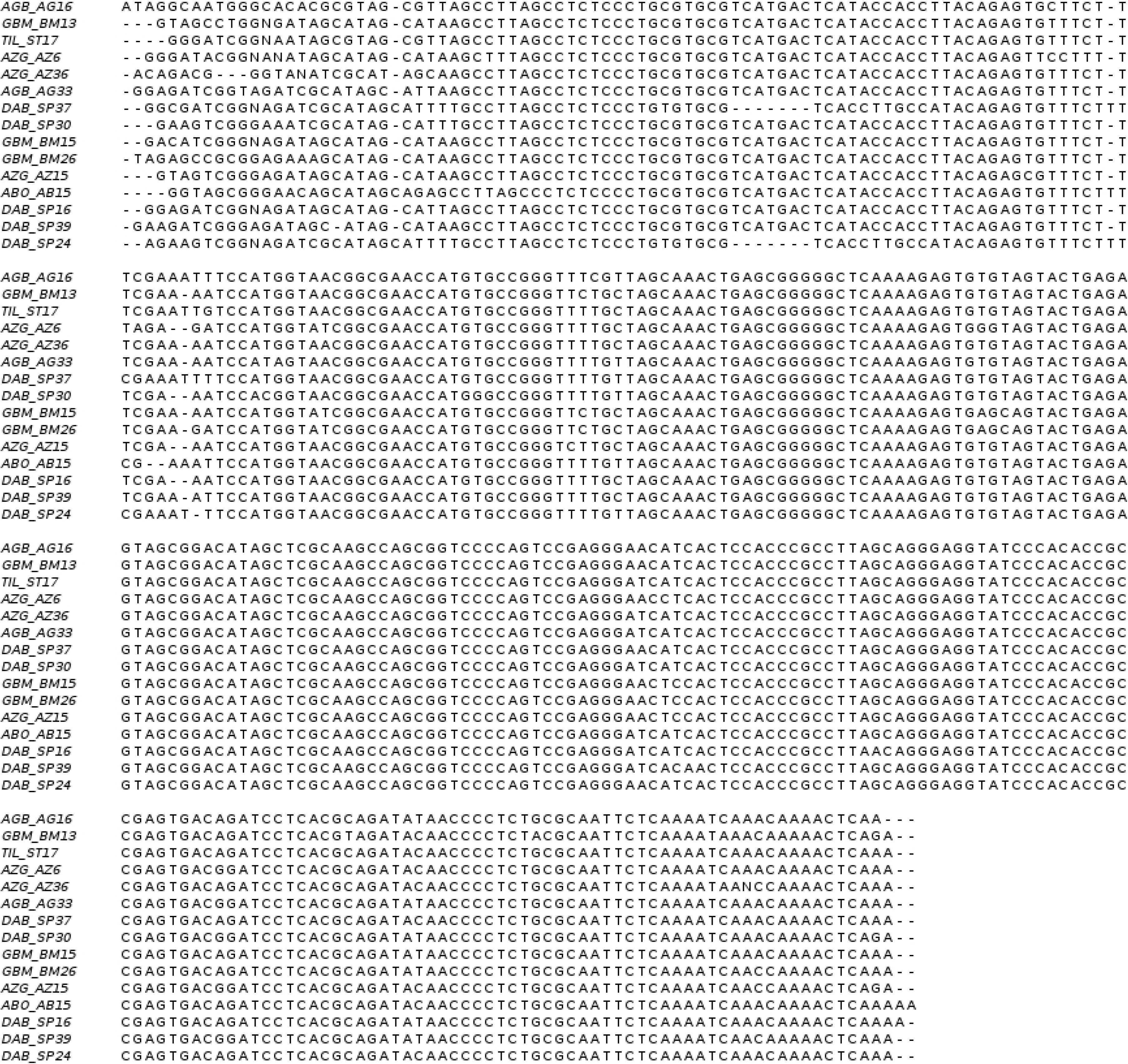
Multiple sequences alignment of satRNAs from Cote d’Ivoire.

**Fig 9.**
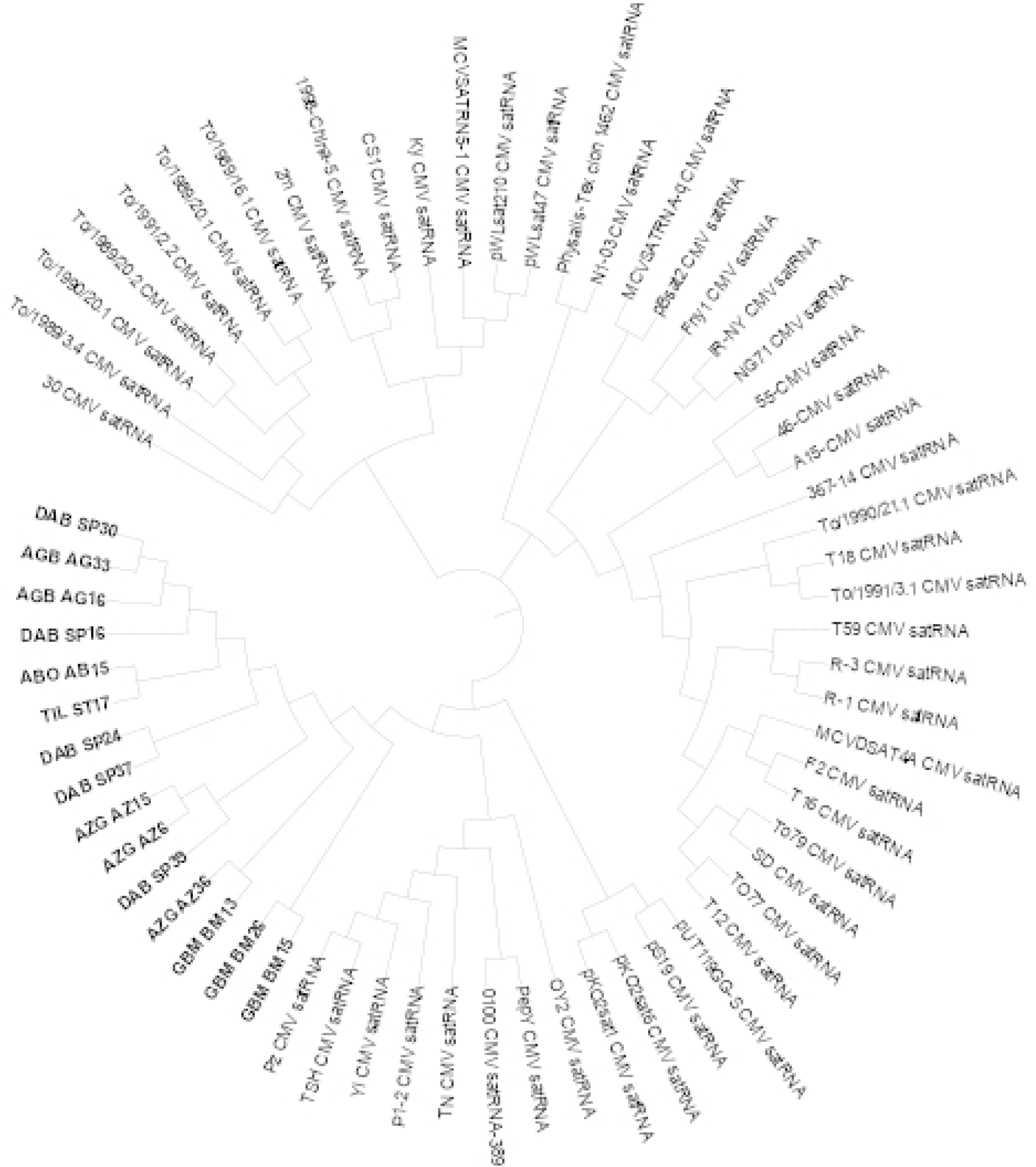
Estimation of Ivorian CMV related satRNAs phylogenetic relationship with known satRNAs using the Maximum Likelihood method and Jukes-Cantor model. The bootstrap consensus tree inferred from 1000 replicates is taken to represent the evolutionary history of the taxa analyzed. Branches corresponding to partitions reproduced in less than 50% bootstrap replicates are collapsed. A Gamma distribution were used to model the substitution rate with 5 discrete Gamma categories.

## Discussion

Côte d’Ivoire is the first African exporter of desert bananas [1]. Our survey aimed to measure the real level of CMV occurrence in industrial banana plantations. A second goal was to achieve a proper molecular characterization of the concerned strains of the virus since symptoms observed on the field are sometimes very confusing. Our study focused on banana varieties including mainly Cavendish, cvs Great Naine and William, AAA group.

We showed that CMV is effectively present in all 7 main production areas of banana in Côte d’Ivoire. Results of our study indicated that virus isolates causing chlorosis, necrosis, and mosaic on *Musa sp.* belonged to subgroup IA (based on molecular analyses). However, symptoms observed on infected leaves varied from plant to plant in a given sampling area and also between areas. We also noticed that these symptoms are uncorrelated with confirmed CMV presence. The latter may suggest the combination of CMVs with other factors like abiotic stress and/or other viruses.

The present study is, to the best of our knowledge, the first report of micro-satellite RNA sequences associated with CMV naturally occurring on *Musa sp.* in Côte d’Ivoire. Further molecular characterization by constructing and analyzing infectious clones will be required for a complete characterization of the impact of these satellites in pathogenicity.

From our molecular characterization, it appeared a high similarity score between coat protein sequences of Ivorian CMVs. In addition to considering this result as good conservation of the concerned strains, we also hypothesize that the observed CMV strains may have the same origin. Indeed, in Côte d’Ivoire, for new banana field installation, farmers actually take clones from their neighborhood, causing the spread of the viral infection from area to area. We, therefore, recommend farmers to pay much attention to banana seeds. Systematic virus detection should be carried out prior to use such seeds to install new fields. In this context, the DAS-ELISA protocol could be proposed since it is economically accessible to fruit companies and will avoid important production losses. Currently, symptomatic plants are detected one month after planting and replaced by healthy ones. However, in some cases, one month is already too late to avoid large contamination. Systematic detection before planting remains the best strategy.

Compared to studies where only the biological indexing is carried out, our results demonstrate the relative difficulty to directly correlate results from biological indexing, serologic data from DAS-ELISA, and genomic information from Sequencing. On the 65 infected samples (confirmed by both DAS-ELISA and Amplicon Sequencing) used to perform biological indexing, only 14 caused symptoms. This demonstrates, once again, the low reliability of biological indexing compared to both DAS-ELISA and amplicon sequencing [10–18].

Based on a single amino acid difference in the coat protein sequences, we distinguished two groups in Ivorian strains. As previously established, it is worth mentioning that a single amino acid variation can induce different symptoms on plants: shifting from light green/dark mosaic to bright yellow/white chlorosis in tobacco [24] or causing white mosaic, pale green mosaic, veinal chlorosis, veinal necrosis, systemic necrosis, and necrotic local lesions and affect thylakoid structure [25]. It is also worth indicating that we only partially sequenced the coat protein. The diversity and somehow inconsistency in symptoms observed during the field survey may reside in additional differences that would appear in the full-length coat protein sequence. This point should be considered for further studies on Ivorian CMV strains infecting *Musa spp* trees. As indicated, we were not able to attribute particular symptoms to specific amino acid variation in Ivorian CMV based on biological indexing. This point needs a deeper analysis along with the presence or absence or micro-satellite. Indeed, the presence of satRNA does not seem to influence pathogenicity: observed symptoms are mainly the same in all collecting places including Aboisso where no satRNA was detected.

Phylogenetic analysis showed that AZ6 (KC189912) sequence and BM26 (KC189911) sequences are closed to E5-CMV and D8-CMV respectively, both coming from Japan. This result can be explained the convergent evolution of CMV strains worldwide.

## Acknowledgments

We thank Prof. Agneroh for guidance, field, and lab work supervision. We thank technicians from the Gembloux Agro-Bio tech lab for guidance and assistance in lab works. We also acknowledge funding received from Gembloux Agro-Bio Tech. Many thanks to fruit companies of Côte d’Ivoire for giving access to their fields for sample collection.

## Supporting information

**S1 Table. Accession number and origin of selected CMV coat protein sequences used for sequence comparison with Ivorian CMVs.** *: IA CMV subgroup, **: IB CMV subgroup, ***: II CMV subgroup

**S2 Table. List of satRNAs sequences used for sequence comparison with Ivorian satRNAs.**

**S3 Table. Symptoms observed on reporting plants after CMV’s biological indexing.** C: Chlorosis, M: Mosaic, FN: Foliar Necrosis, CR: Cigar Rot, LD: Leaf Distortion, LC: Leaf Curling, D: Dwarfing of the banana tree, TS: Thin Sheet

## References

1. FAOSTAT. [cited Sep 3rd 2020]. Available: http://www.fao.org/faostat/en/#rankings/commodities_by_country

2. ABIDJAN.NET [Online Press article published on Dec 3rd 2019]. Banane dessert: la Côte d’Ivoire, 1er producteur africain avec près de 450 000 tonnes en 2019. Available at https://news.abidjan.net/h/666652.html

3. Scholthof K-BG, Adkins S, Czosnek H, Palukaitis P, Jacquot E, Hohn T, et al. Top 10 plant viruses in molecular plant pathology. Mol Plant Pathol. 2011;12: 938–954. doi:10.1111/j.1364-3703.2011.00752.x

4. Jacquemond M. Cucumber mosaic virus. Adv Virus Res. 2012;84: 439–504. doi:10.1016/B978-0-12-394314-9.00013-0

5. Kim M-K, Kwak H-R, Lee S-H, Kim J-S, Kim K-H, Cha B-J, et al. Characteristics of Cucumber mosaic virus isolated from Zea mays in Korea. Plant Pathol J. 2011;27: 372–377. doi:10.5423/PPJ.2011.27.4.372

6. Roossinck MJ, Zhang L, Hellwald K-H. Rearrangements in the 5′ Nontranslated Region and Phylogenetic Analyses of Cucumber Mosaic Virus RNA 3 Indicate Radial Evolution of Three Subgroups. J Virol. 1999;73: 6752–6758.

7. Qiu Y, Zhang Y, Wang C, Lei R, Wu Y, Li X, et al. Cucumber mosaic virus coat protein induces the development of chlorotic symptoms through interacting with the chloroplast ferredoxin I protein. Sci Rep. 2018;8: 1205. doi:10.1038/s41598-018-19525-5

8. Owen J, Shintaku M, Aeschleman P, Tahar SB, Palukaitis P. Nucleotide sequence and evolutionary relationships of cucumber mosaic virus (CMV) strains: CMV RNA 3. J Gen Virol. 1990;71: 2243–2249. doi:10.1099/0022-1317-71-10-2243

9. Lin H-X, Rubio L, Smythe A, Jiminez M, Falk BW. Genetic diversity and biological variation among California isolates of Cucumber mosaic virus. J Gen Virol. 2003;84: 249–258. doi:10.1099/vir.0.18673-0

10. Dubey VK, Aminuddin, Singh VP. Molecular characterization of Cucumber mosaic virus infecting Gladiolus, revealing its phylogeny distinct from the Indian isolate and alike the Fny strain of CMV. Virus Genes. 2010;41: 126–134. doi:10.1007/s11262-010-0483-6

11. Hsu Y-H, Wu C-W, Lin B-Y, Chen H-Y, Lee M-F, Tsai C-H. Complete genomic RNA sequences of cucumber mosaic virus strain NT9 from Taiwan. Arch Virol. 1995;140: 1841–1847. doi:10.1007/BF01384346

12. Hord MJ, García A, Villalobos H, Rivera C, Macaya G, Roossinck MJ. Field Survey of Cucumber mosaic virus Subgroups I and II in Crop Plants in Costa Rica. Plant Dis. 2001;85: 952–954. doi:10.1094/PDIS.2001.85.9.952

13. Aka AR, Kouassi NK, Agnéroh TA, Amancho NA, Sangare A. Distribution et incidence de la mosaïque du concombre (cmv) dans des bananeraies industrielles au sud-est de la côte d’ivoire. Sci Nat. 2009;6. doi:10.4314/scinat.v6i2.48670

14. Ayo-John EI, d’Arros HJ, Ekpo EJA, Shoyinka SA. A survey in Southern Nigeria reveals the presence of *Cucumber mosaic virus* subgroup I in *Musa* crops. Fruits. 2008;63: 135–143. doi:10.1051/fruits:2008003

15. Sutula C, Gillet J, Morrissey S, Ramsdell D. Interpreting ELISA data and establishing the positive-negative threshold. Plant Dis. 1986;70: 722–726.

16. Sharman M, Thomas JE, Dietzgen RG. Development of a multiplex immunocapture PCR with colourimetric detection for viruses of banana. J Virol Methods. 2000;89: 75–88. doi:10.1016/S0166-0934(00)00204-4

17. Bariana HS, Shannon AL, Chu WG, Waterhouse PM. Detection of five seedborne legume viruses in one sensitive multiplex polymerase chain reaction test. Phytopathology. 1994;84: 1201–1205.

18. Gafny R, Wexler A, Mawassi M, Israeli Y, Bar-Joseph M. Natural infection of banana by a satellite-containing strain of cucumber mosaic virus: nucleotide sequence of the coat protein gene and the satellite RNA. Phytoparasitica. 1996;24: 49–56. doi:10.1007/BF02981453

19. Nouri S, Arevalo R, Falk BW, Groves RL. Genetic Structure and Molecular Variability of Cucumber mosaic virus Isolates in the United States. PLOS ONE. 2014;9: e96582. doi:10.1371/journal.pone.0096582

20. Sievers F, Wilm A, Dineen D, Gibson TJ, Karplus K, Li W, et al. Fast, scalable generation of high-quality protein multiple sequence alignments using Clustal Omega. Mol Syst Biol. 2011;7: 539. doi:10.1038/msb.2011.75

21. Camacho C, Coulouris G, Avagyan V, Ma N, Papadopoulos J, Bealer K, et al. BLAST+: architecture and applications. BMC Bioinformatics. 2009;10: 421. doi:10.1186/1471-2105-10-421

22. Kumar S, Stecher G, Li M, Knyaz C, Tamura K. MEGA X: Molecular Evolutionary Genetics Analysis across Computing Platforms. Mol Biol Evol. 2018;35: 1547–1549. doi:10.1093/molbev/msy096

23. Letunic I, Bork P. Interactive Tree Of Life (iTOL) v4: recent updates and new developments. Nucleic Acids Res. 2019;47: W256–W259. doi:10.1093/nar/gkz239

24. Shintaku MH, Zhang L, Palukaitis P. A single amino acid substitution in the coat protein of cucumber mosaic virus induces chlorosis in tobacco. Plant Cell. 1992;4: 751–757. doi:10.1105/tpc.4.7.751

25. Mochizuki T, Ohki ST. Single amino acid substitutions at residue 129 in the coat protein of cucumber mosaic virus affect symptom expression and thylakoid structure. Arch Virol. 2011;156: 881–886. doi:10.1007/s00705-010-0910-y

